# RNA beacons for detecting the human testis-determining factor

**DOI:** 10.1101/2021.04.22.440928

**Authors:** Diego F. Joseph, Jose Alberto Nakamoto, Pohl Milón

## Abstract

The testis-determining factor (TDF) is an essential transcriptional protein for male differentiation in mammals, expressed along spermatids to early zygotes and, to some extent, in diverse cellular lines. In this study, we developed fluorescent biosensors capable of indicating the presence of TDF. We used *in vitro* evolution techniques to produce RNA aptamers that bind the recombinantly expressed HMG-box, the DNA binding domain of TDF. Bioinformatic analysis along *in vitro* evolution setup suggested two predominant aptamer clusters with distinctive motifs. The top ranked aptamer from each cluster, M1 and M2, showed specific binding to TDF. Aptamers were fluorescently modified as molecular beacons. Pre-steady-state kinetics indicated the beacons bind rapidly, within 50 seconds, yet M1 showed better signal to noise ratios than M2. Structural predictions of the aptamer interaction indicated that M1 is composed by three stem loops and likely interact with the HMG-box of TDF through the pocket formed by the three loops. Molecular modelling of M1 beacon shows that binding to TDF entails a conformational change of the sensor resulting in the measured fluorescence changes. To our knowledge, this is the first work describing an RNA beacon for detecting the essential TDF. Potential applications and advantages over alternative methods are provided and discussed.

## 1. Introduction

Sexual differentiation is established since fertilization; particularly, in mammals, the presence of a normal Y chromosome coordinates the development of a male phenotype (Goodfellow and Darling 1988). Male differentiation is due to a gene located on the short branch of the Y chromosome called the sex-determining region Y or so called *SRY* gene (George and Wilson 1984). The *SRY* gene is intronless and code for a transcript of 1.1 kilobases which is then translated into a protein of 204 amino acids with a molecular mass of 23.9 KDa called the testis-determining factor (TDF) (Ellis and Erickson 2017; Su and Lau 1993).

TDF can be divided into three main regions: The C-terminal domain, which has no known conserved structure; the N-terminal domain, which can be phosphorylated to enhance DNA binding; and the central region, which has a conserved DNA binding domain: the high mobility group box or so called HMG-box (Denny et al. 1992; Harley, Clarkson, and Argentaro 2003; Sinclair et al. 1990). TDF, along other proteins that share the HMG-box domain, are part of a gene family called the SRY-related box (*SOX*) (Kiefer 2007). These proteins play a crucial roles as transcription factors during embryo development, being TDF and SOX9, the male determination factors (Harley, Clarkson, and Argentaro 2003; Kanai et al. 2005; Kiefer 2007).

Since the Y chromosome is exclusively found in normal mammalian males; the presence of both, the *SRY* transcript and TDF are limited to male mammalian cells. In the case of the *SRY* transcript, it has been reported to be present in human cell lines such as DU145 and HepG2, as well as in different human tissues (Clepet et al. 1993; Tricoli et al. 1993). Moreover, different publications have reported the presence of the *SRY* transcript in the human Sertoli and germ cells, as well as in ejaculated human Y-spermatozoa (Modi et al. 2005; Salas-Cortés et al. 1999, 2001). Regarding its localization, Modi *et al*. reported the presence of the *SRY* transcript exclusively in the midpiece of ejaculated human Y-spermatozoa (Modi et al. 2005). In the case of the TDF, it has been reported to be present and localized in the nuclei of human cells lines such as NT2D1, HS68, and in *SRY*-transfected HeLa cells, as well as Sertolli cells and round spermatids (Poulat et al. 1995; Salas-Cortés et al. 1999, 2001). No conclusive data could be found about the localization of the TDF in human ejaculated spermatozoa. Nonetheless, Li *et al*. reported in 2011 that the TDF is found in the head of mature bovine Y-spermatozoa (Li et al. 2011). Altogether, TDF is an essencial transcriptional factor for male differenciation across mammals, especifically expressed in male cell lines and male spermatis and therefore a potential biomarker for the development of novel methods for early sexual identification.

In this work, we aimed to develop RNA aptamers as specific binders of TDF and sought to bioengineerized them into molecular beacons for TDF detection (Figure 1A). Aptamers are nucleic acid molecules that bind with high affinity to a specific target molecule (Tuerk and Gold 1990; Wu and Kwon 2016). Aptamers offer a tentative alternative for the development of biosensors as their development do not require immunogenic molecules (Thiviyanathan and Gorenstein 2012). Moreover, as they are obtained by affinity through the Systematic Evolution of Ligands by Exponential enrichment (SELEX) (R. Oliphant, J. Brandl, and Struhl 1989; Tuerk and Gold 1990), they do not compromise the binding and biochemical specificity parameters (Thiviyanathan and Gorenstein 2012). Furthermore, aptamers can be modified with fluorophores and work as beacons when binding to their target molecule (Hamaguchi, Ellington, and Stanton 2001).

**Figure 1.**
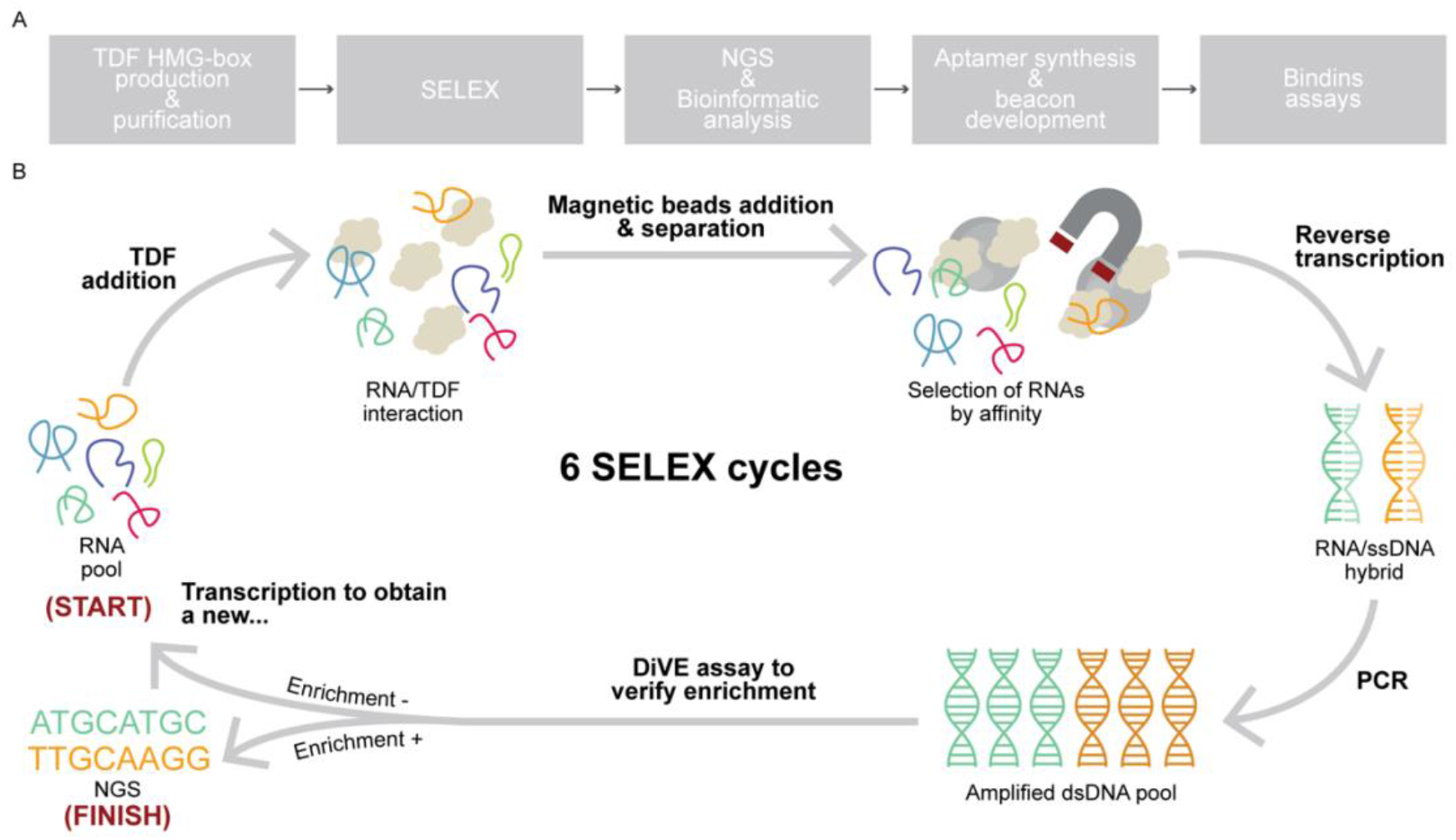
Experimental approach for RNA aptamer selection and characterization. (A) General workflow for developing RNA aptamers for the TDF. (B) Scheme of the followed SELEX method: The RNA library (N30) was incubated with the recombinant and his-tagged TDF protein. After incubation, Ni-NTA-modified magnetic beads were added and were incubated for a short period. Then, a magnet was used to separate the RNA aptamers-TDF-magnetic beads complexes from the non-bound oligonucleotides. In order to amplify the bound oligonucleotides, *in vitro* reverse transcription step was needed to obtain an RNA/DNA hybrid. Then, a PCR step was performed to obtain a dsDNA. After every PCR step, a DiVE assay was carried out to check the enrichment of the oligonucleotides pool. If the enrichment was reached, the samples were processed for Next-Generation-Sequencing (NGS); otherwise, the T7 *in vitro* transcription step was performed in order to proceed to a new SELEX cycle.

Thus, aptamers can provide alternatives to the reported antibodies for the detection or localization of the human TDF (Kodrič et al. 2019; Olivares et al. 2018; Salas-Cortés et al. 1999, 2001). Here, we use recombinant biology to produce and purify the HMG-box domain of the human TDF as target for SELEX to develop high-affinity RNA aptamers, and further modify them into molecular beacons to detect the human TDF.

## 2. Materials and methods

### 2.1 Cloning, expression, and purification of the TDF HMG-box

The human TDF HMG-box (hereinafter called TDF) was cloned into the plasmid pET-24c (+) (KanR) by GenScript (USA) using the sequence from the position 313 to 528 of the mRNA. The coding sequences were optimized for codon usage of *E. coli* BL21 using OptimumGene^™^ (GenScript, USA) and a His-Tag was added at the carboxyl end. Competent *E. coli* were produced using Mix & Go *E. coli* Transformation Kit & Buffer Set (Zymo Research, USA) and used for bacterial transformation. Transformed cells were grown in Luria-Bertani broth (LB) (tryptone 10g, NaCl 10g, yeast extract 5g, H2O to 1L) at 37°C and continuous shaking until absorbance at 600 nm reached 0.5 O.D. Then, 1 mM isopropyl β-D-1-thiogalactopyranoside was added to induce protein expression and incubation continued for 4 hours. Cells were pelleted by centrifugation at 4,000 x g for 10 min at 4°C and the supernatant was discarded. Cell pellets were washed by resuspending them in HAKM10 buffer (50 mM HEPES pH 7.4, 70 mM ammonium acetate, 30 mM NaCl, 10 mM MgCl2, 6 mM 2-mercaptoethanol). The cells were centrifuged at 6 000 x g for 20 min at 4°C and the supernatant was discarded. Cells were resuspended in 5 mL of His-tag Buffer (20 mM sodium phosphate pH 7.4, 0.5 M NaCl) + 10 mM imidazole mL for each gram of cells and lysed by sonication. Then, the lysate was centrifuged at 10 000 x g for 45 min at 4°C. The supernatant was then collected for purification using a 5 mL His Trap^™^ FF Crude Column containing Ni-NTA (GE Healthcare, USA). The column was first equilibrated using five volumes of His-tag Buffer + 10 mM imidazole and the supernatant was loaded into de column. The flow-through was discarded. After that, 10 volumes of His-tag Buffer + 10 mM imidazole followed by 10 volumes of His-tag Buffer + 50 mM imidazole were passed through the column to elute nonspecifically bound proteins. Finally, 5 mL of His-tag Buffer + 500 mM imidazole were used to elute the protein. Fractions of 0.5 mL were collected. The purest and most concentrated fractions were then dialyzed by employing a 3 kDa membrane D-Tube Dialyzer (Merck, Germany) in PBS buffer (200 mM NaCl, 2.7 mM KCl, 10 mM Na2HPO4, 1.8 mM KH2PO4, 10% glycerol, pH 7.4). Finally, TDF purity and concentration were determined by (18% acrylamide) SDS-PAGE electrophoresis and Coomassie blue staining and by Bradford Protein Assay.

### 2.2 RNA library and SELEX workflow

A N30 RNA random library (Trilink Biotechnologies, USA) was used for the SELEX procedure. Each library contains the following features: RNA Template (5’ UAG GGA AGA GAA GGA CAU AUG AU (N30) UU GAC UAG UAC AUG ACC ACU UGA 3’), T7-Promoter Forward Selection Primer (5’ TTC AGG TAA TAC GAC TCA CTA TAG GGA AGA GAA GGA CAT ATG AT 3’), and Reverse Selection Primer (5’ TCA AGT GGT CAT GTA CTA GTC AA 3’). Each SELEX cycle included an TDF-RNA pool incubation, Ni-NTA magnetic beads addition and magnetic separation, reverse transcription of the TDF-bound RNAs, PCR, and transcription. Each cycle also had a previous incubation of the RNA pool with the human HMG-box of the High Mobility Group Box 1 as a negative control. Figure 1B shows a detailed scheme of the SELEX cycle. Purity and size of the double stranded DNA (dsDNA) were assessed by native PAGE 8%, while for RNA were assessed by 8% 8M urea PAGE. Nucleic acids concentration was measured by absorbance at 260 nm using a Nanodrop One spectrophotometer (Thermo Fisher Scientific, USA). All selections were performed using SELEX buffer (100 mM NaCl, 1.5 mM KCl, 5 mM MgCl2, 50 mM Tris HCl, pH 7.4) at 25°C and the stringency was increased along cycles by modifying the ionic strength, time of incubations and number of washes. Refer to Supplementary Table 1 for more details.

### 2.3 S1 endonuclease as SELEX pool enrichment indicator

The S1 endonuclease hydrolyses single-stranded nucleic acids structures such as ends, nicks, and heteroduplex loops into oligo or mononucleotides. This ability let us differentiate between enriched and non-enriched SELEX pools of potential aptamers, being the enriched ones the less degraded. This assay is better known as the Diversity Visualization by Endonuclease (DiVE) assay and it was followed as described by *Lim et al* (Lim et al. 2011). For this assay, we prepared seven reaction tubes (one for each SELEX cycle + one control: enriched homoduplex) of 200 ng of dsDNA in 20 μL of S1 reaction buffer 1X (40 mM CH3COONa, 300 mM NaCl, 2 mM ZnSO4, pH 4.5). After 5 min of denaturation at 95°C and 5 min of stringent renaturation at 65°C, 1 U/μL of S1 endonuclease was added and then incubated at 65°C for 30 min. Finally, the reactions were stopped using EDTA into a final concentration of 2 mM. Approximately, 50 ng of each sample was mixed with RunSafe (Cleaver Scientific U.K.) and visualize in a 2% agarose gel.

### 2.4 Next-Generation-Sequencing

For sequencing, the dsDNA from each of the six SELEX cycles was used. These dsDNA pools were processed as described in the 16S Metagenomic Sequencing Library preparation protocol (Illumina, USA) (Illumina 2013). In summary, 0.1 pmol dsDNA was used to perform the 1^st^ stage PCR in which forward and reverse primer overhang adapters were employed. Then, the first PCR clean-up was performed using Ampure XP Beads (Beckman Coulter, Inc) and 80% ethanol. After the clean-up, the 2^nd^ PCR stage was performed using the Nextera XT Index 1 and Index 2 primers from the Nextera XT Index Kit Illumina (California, USA). Then, the second PCR clean-up was performed using the same procedure as described above. After that, the pools were quantified, normalized and mixed together. Finally, the combined samples were diluted and denatured using 0.2 N NaOH, mixed with the PhiX Control Kit V3, and were sequenced using the MiSeq Reagent Nano Kit v2 cartridge (Illumina, USA) for 300 reading cycles.

### 2.5 Bioinformatic analysis and aptamer ranking

Retrieved data from sequencing was previously processed in order to be analyzed. The data corresponding to the reverse strand was reverse complemented using the QC and Reversecomplement functions from the Galaxy Project platform and then added to the forward data (Afgan et al. 2018). After that, the whole new data was trimmed to remove the conserved primer flanking regions using the QC and Trim Sequences functions from the Galaxy Project platform (Afgan et al. 2018).

A primary analysis of the sequences and a comparison between SELEX pools were performed using the bioinformatic toolkit FASTAptamer (Alam, Chang, and Burke 2015) essentially as described by Joseph *et al*. (Joseph et al. 2019). First, the FASTAptamer Count command was used to estimate the frequency of the sequences in each SELEX cycle. Then, the FASTAptamer Cluster command was employed using a Levenshtein edit distance of 7 and sequences with more than 5 RPM were used to identify aptamer families. Data obtained from the FASTAptamer Cluster analysis was used to identify the most ranked aptamers by family using the FASTAptamer Enrich command, and to identify motifs using The MEME Suite (Bailey et al. 2009). Finally, Mfold was used to determine Gibbs free energy of the most ranked sequences obtained (Zuker 2003).

### 2.6 3D modelling and docking predictions

The 3D structure of TDF was built through homology modelling using the SWISS-MODEL suite (Waterhouse et al. 2018). The 3D structure for the complete TDF was built through threading using the PHYRE 2 Protein Fold Recognition Server (Kelley et al. 2016). Aptamer secondary structures were predicted using the RNAfold web server and output files were saved in the Vienna bracket format (Gruber et al. 2008). Single stranded RNA 3D models were built using the RNAComposer server (Antczak et al. 2016; Popenda et al. 2012a). The aptamer structure was protonized simulating the SELEX conditions (pH: 7.4, 100 mM salt concentration) and subsequently refined through energy minimization using MOE software (MOE 2020). Rigid structure docking of the TDF to the aptamers was perform using MOE software (MOE 2020).

### 2.7 Oligonucleotides synthesis and annealing

The top ranked aptamer of each of the two most representative motifs obtained by the previous analysis, as well as the probes that were employed for the following assays, were synthesized by Macrogen (Korea). Refer to Supplementary Table 2 for more details about the sequences. The two dsDNA were used for *in vitro* transcription following the protocol stablished by the TranscriptAid T7 High Yield Transcription Kit (Thermo Fisher Scientific, USA). Because our transcripts were under 100 nucleotides, 2 μg of dsDNA template were used instead of 1 μg. The reactions were incubated at 37°C for 2.5 hours. After incubation, 4 U of RNAse-free DNAse I were added to each reaction tube and incubated for 15 min at 37°C to remove the DNA template. Finally, our transcripts were purified using the Quick-RNA Miniprep Kit (Zymo Research, USA) and measured by absorbance at 260 nm using a Nanodrop One spectrophotometer (Thermo Fisher Scientific, USA). For the probes and aptamer annealing, equal concentrations of probes and aptamer were mixed and incubated for 3 min at 70°C and then incubated at RT for at least 15 min prior use.

### 2.8 TDF pull down

Our two top ranked aptamers (named M1 and M2) were subjected to interact with the TDF in order to determine their binding specificity. Briefly, seven reaction of 500 μL (SELEX buffer, 0.02% Tween^™^ 20) were prepared; each containing 100 nM of protein (TDF or control) and 300 nM of oligonucleotides (M1, M2 or control). Each reaction was made in triplicates. The unrelated PKnG kinase domain of *Mycobacterium tuberculosis*, and no-bound SELEX 1 library were used as controls. After 20 min of incubation, Ni-NTA coated magnetic beads (Dynabeads^™^ His-Tag Isolation and Pulldown from Invitrogen, USA) were added to the solution and incubated for 10 additional min in order to bind the his-tagged TDF and PKnG proteins. After that, magnetic beads, bound proteins, and attached aptamers were separated using a magnet. Then, the magnetic beads were resuspended in 20 μL of SELEX buffer 1X and incubated at 95°C for 3 min. Finally, the supernatant was obtained and the nucleic acids in there were measured by absorbance at 260 nm using a Nanodrop One spectrophotometer (Thermo Fisher Scientific, USA).

### 2.9 Kinetic experiments

An SX20 stopped-flow apparatus (Applied Photophysics, UK) was used to measure the interactions of our aptamers M1 and M2 and the TDF in real time. All reactions were performed using SELEX buffer pH 7.4 as the solution for aptamer/protein interaction. All fluorescent measurements were carried out at 25 °C using a monochromatic light source of 470 nm powered with 10 mA while the emitted fluorescence was measured using photomultipliers previously set at 335 V after passing a long-pass filter with a cutoff 515 nm (Chulluncuy et al. 2016). Each reaction measured 7 replicates, and every measurement acquired 1000 data points during 150 seconds in a logarithmic sampling mode.

### 2.10 Determination of kinetic constants

Non-linear regression with exponential equations were used to estimate the apparent constants. All the equations and graphics were made using Prism 9 (Graphpad Sofware, USA). Averaged rate constants were calculated as described by Milon *et al*. (Milon et al. 2008).

## 3. Results

### 3.1 Aptamers against the human TDF

The Systematic Evolution of Ligands by Exponential enrichment (SELEX) method, combined with the Next-Generation Sequencing (NGS), allowed us to develop molecular sensors that can specifically recognize the human TDF. The SELEX method permits an enrichment of singlestranded oligonucleotides that bind with high affinity a given molecule (Tuerk and Gold 1990). These oligonucleotides, named aptamers, have shown to have significant advantages such as thermal stability and being able to recognize any kind of molecule in addition to immunogenic proteins (Stoltenburg, Reinemann, and Strehlitz 2005; Thiviyanathan and Gorenstein 2012). These features seem ideal to develop aptamers to detect a protein that is exclusively expressed male cells, and even in germ cells such as round spermatids (Salas-Cortés et al. 1999, 2001). The SELEX method was based as described by Joseph *et al*. (Joseph et al. 2019). We performed six SELEX cycles starting from a library of RNA containing 30 randomized nucleotides (Figure 1B). To enhance the selection of positive binders, the concentration of the protein, incubation time, and number and strength of washes varied (Supplementary Table 1). As for increasing the specificity of our potential aptamers, we performed negative selections in each cycle using the human HMG-box of HMGB1, a conserved domain along all transcription factors. Since the input RNA pool from each RNA/TDF interaction is very low, we did not measure the RNA output concentration, but proceed directly with reverse transcription and PCR. The dsDNA product was measured; however, we did not find any correlation between dsDNA concentration and SELEX cycle (Supplementary Table 1). No correlation was either observed when performing *in vitro* transcription, despite this step had varying amounts of starting dsDNA.

During SELEX cycles, it is expected a loss of variability while enriching a sub-population of potential aptamers. This enrichment is assed using the DiVE assay and a bioinformatic analysis (Lim et al. 2011; Schneider, Rasband, and Eliceiri 2012). While we used an RNA library, there is a step where dsDNA is generated in order to amplify our library, a step that is suitable for the DiVE assay. DiVE uses the S1 endonuclease to differentiate between dsDNA homoduplexes and heteroduplexes by cleaving single stranded areas found in heteroduplexes. Thus, an ideal enriched pool will be visualized as a sharp band in an agarose gel, without being cleaved. The DiVE assay allowed us to confirm enrichment along all the SELEX cycles. There were no bands in samples of cycles 1 and 2, which reflects that all the reannealed dsDNA was cleaved and there was no enrichment of sequences in these pools. In samples 3 and 4, slight bands are shown; while these bands become stronger in samples 5 and 6, which confirms that some sequences are starting to get enriched (Supplementary Figure 1). All SELEX cycles were used for NGS using the Illumina system (Illumina 2013).

In order to analyze the NGS output, we followed the bioinformatic workflow described in figure 2A. The outputs from the Galaxy Project (see 2.5 Bioinformatic analysis and aptamer ranking) were processed using the FASTAptamer-Count algorithm of the FASTAptamer toolkit (Alam, Chang, and Burke 2015). This algorithm ranks the sequences based on their abundance in each file, which represents SELEX cycles. Each cycle had in average 96500 individual reads, varying from 32900 to 131600 individual reads (See raw ranked sequences in Supplementary Data 1). The top 5 ranked sequences in cycle 6 varied from 3765 Reads Per Million (RPM) to 32943 RPM, while in cycle 1 varied from 939.01 RPM to 11920.85 RPM. Cycle 6 provided fewer reads, but with a higher frequency, while cycles 1 to 5 showed mostly sequences with less than 12000 RPM of representation (Supplementary Data 2). FASTAptamer-Count algorithm additionally shows the number of unique sequences and the number of total sequences in each SELEX cycle. From this information, we generated a ratio of unique to total sequences to generate an indicator of the global enrichment along the SELEX cycles. A clear enrichment is observed starting from cycle 4 (Figure 2B).

**Figure 2.**
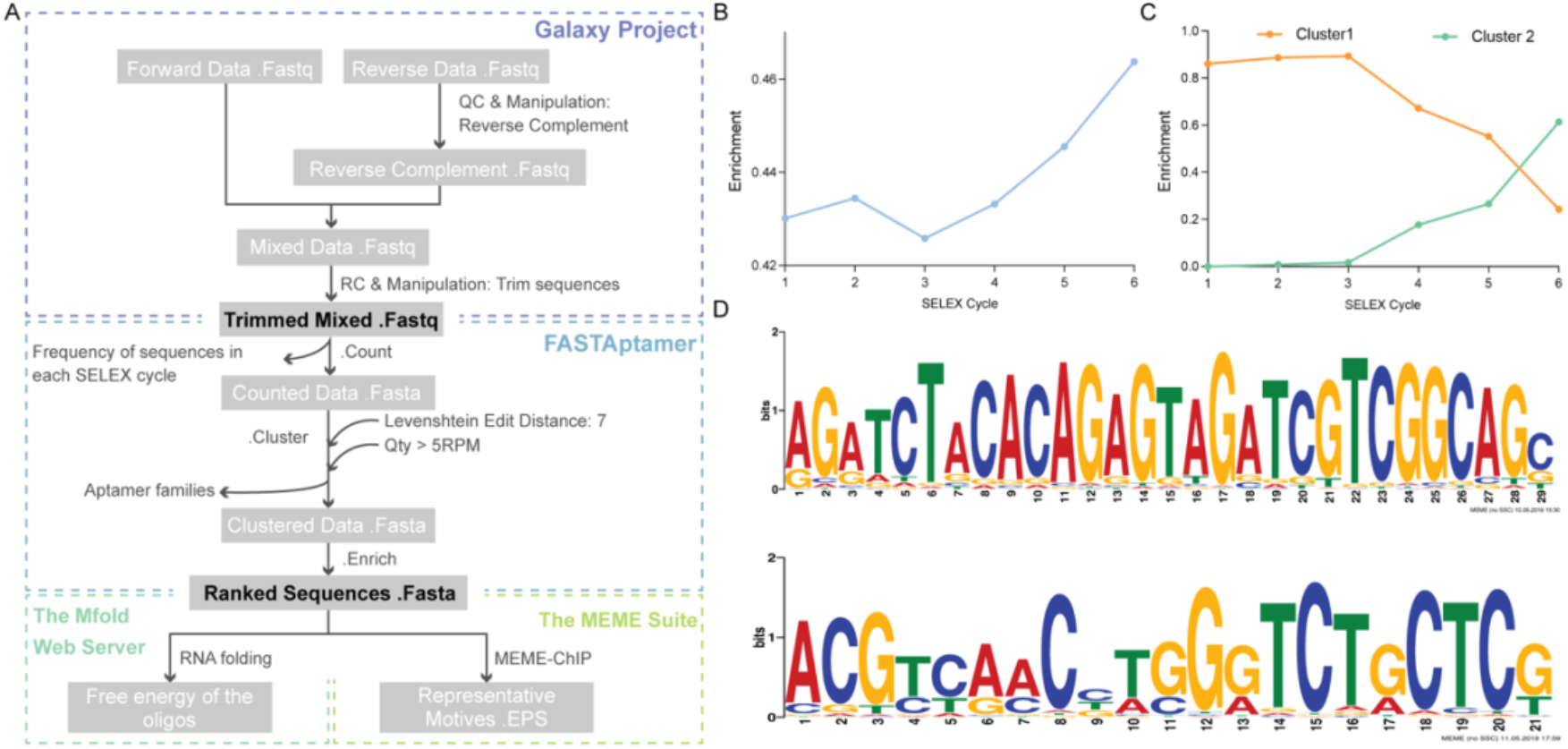
NGS along the bioinformatic toolkits FASTAptamer and the MEME Suite allow the identification of aptamer motifs and their evolution during SELEX cycles. (A) Scheme of the detailed workflow for sequence analysis. (B) Progression of enrichment along the SELEX cycles evaluated by FASTAptamer-Count algorithm. (C) Progression of enrichment of the two representative clusters along the SELEX cycles evaluated by FASTAptamer-Cluster algorithm. (D) The two most representative motifs from SELEX cycle 6 using the MEME-chIP tool from the The MEME Suite webserver.

The above data was used to generate clusters of closely related sequences using the algorithm FASTAptamer-Cluster with a Levenshtein distance of 7 followed by FASTAptamer-Enrich to estimate the ratio of frequencies of each cluster across SELEX cycles. We compared the frequencies of sequences and clusters among cycles 1, 2, & 3 and cycles 4, 5, & 6. Two prominent clusters were identified among the six SELEX cycles. Figure 2C shows the enrichment of both clusters along the SELEX cycles. Cluster 1 had a relative abundance of over 80% from cycle 1 to 3, but then lost representativeness from cycle 4 to 6. On the other hand, the cluster 2 was almost absent until cycle 4 where it started gaining relative abundance up to 61%. The top ranked sequence of the clusters 1 and 2 showed a frequency ratio of 4.68 and 0.89, respectively; in cycles six over five. Moreover, the top ranked sequence of cluster 2 showed a frequency ratio of 0.84 in cycles six over one. Likely due to the loss of representation in late selection. The ratio between cycles six and one could not be calculated for the top ranked family since it was nearly absent in cycle 1 (Supplementary Data 3). Both clusters were analyzed using the the MEME-chIP tool to define the most representative motifs in each cluster. Cluster 1 showed a representative motif of 29 nucleotides, while cluster 2 showed a representative motif of 21 nucleotides (Figure 2D). The top ranked sequence (which includes the representative motif of its cluster) of each of the two clusters were structurally analyzed by the Mfold web server. Since the SELEX cycles were made using not only the variable N30 region, but also the constant regions; the secondary structure and all the incoming experiments were made using the entire sequence: 5’ (N23 constant) N30 variable (N23 constant) 3’. Folding of our two sequences, hereinafter called M1 and M2 aptamers, had a Gibbs free energy of −34.69 Kcal/mol and −26.34 Kcal/mol, respectively.

### 3.2 Structural approximation of the M1/M2–TDF complex

To evaluate the theoretical interaction between the TDF and the M1 or M2 aptamer, we generated 3D models of the TDF domain and the aptamers and used them in a molecular docking approximation. Figure 3A shows the followed workflow to get the molecular docking approximation. In brief, the aptamers secondary structures were predicted using Mfold Web 10 Server (Zuker 2003). Then, 3D models were generated through the RNAComposer tool (Popenda et al. 2012b). The resulting models were refined through energy minimization using MOE software (MOE 2020). M1 predictions build a triple stem loop structure with a small number of unpaired nucleotides, while M2 resulted in a triple stem loop with helix structures between loops 1 and 2 (Supplementary Figure 2).

**Figure 3.**
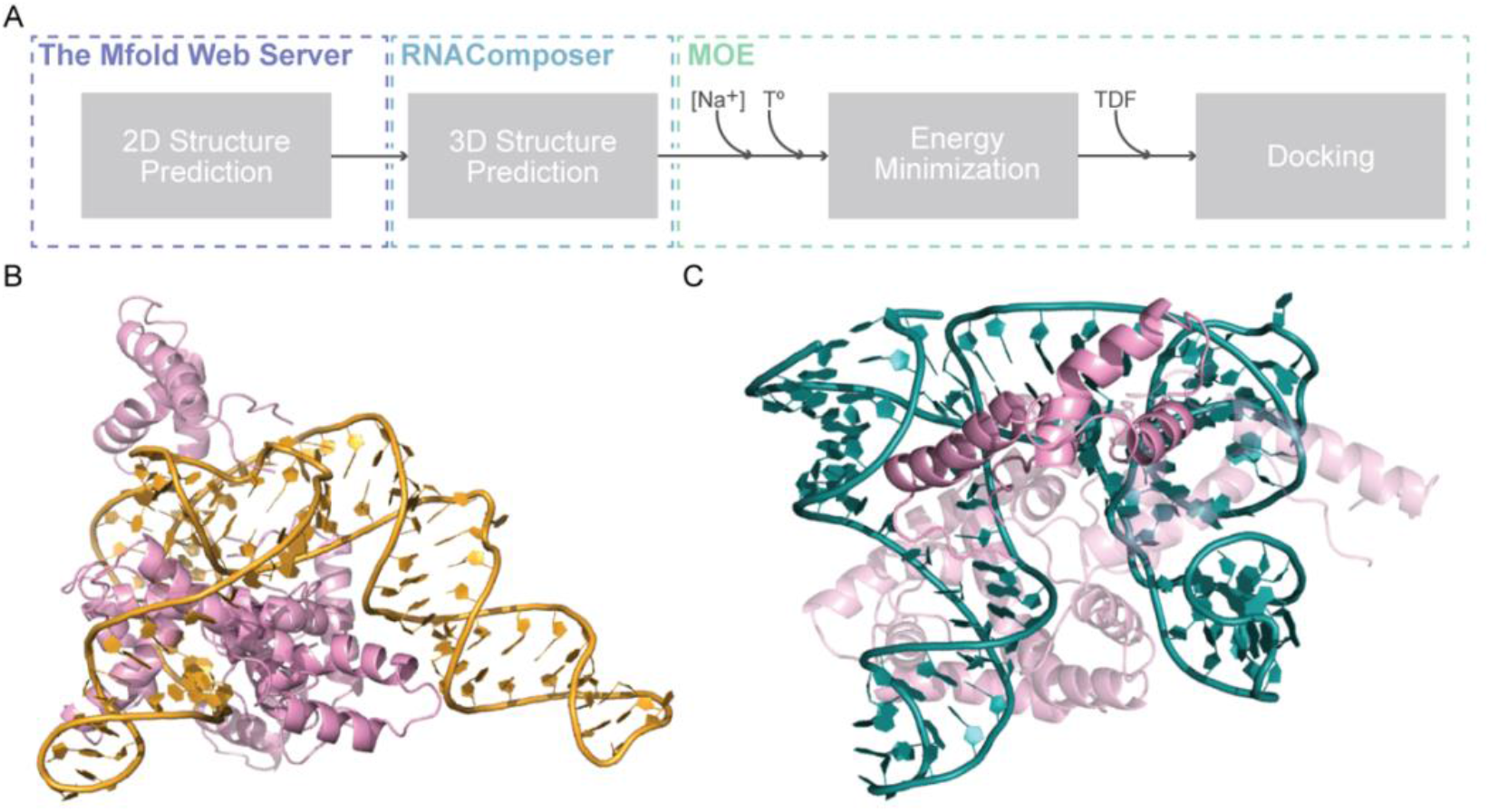
Workflow for the structural approximation of the M1/M2–TDF complex and M1/M2–TDF docking. (A) Scheme of the detailed workflow for the structural approximation to the M1/M2-TDF complex. (B) Structural models of the top five M1-TDF complexes. In the docking with the highest S-score, TDF is shown in a solid opaque color while in the following top S-scores TDF loses opacity. (C) Structural models of the top five M2-TDF complexes. In the docking with the highest S-score, TDF is shown in a solid opaque color while in the following top S-scores TDF loses opacity.

The TDF 3D structure was built through homology modelling with the PDB: 1J46.1 as template by the SWISS-MODEL suite (Waterhouse et al. 2018). For the complete TDF protein, modelling by threading with PHYRE2 resulted in a 3D model with highly unstructured regions apart from the HMG-box domain and was deemed unfit for molecular docking (Supplementary Figure 3). Thus, we used the HMG-box for further structural modeling, from here in called TDF. Modeling of the TDF-Aptamer complexes was performed using the predicted tertiary structures of the aptamer and the protein (see above) with the MOE software (MOE 2020). One hundred complexes for each aptamer were generated. The average value of the S-scores did not indicate substantial differences between the M1 and M2 aptamers. While the top ranked S score for M1 and M2 were −86 and −74 respectively. M1 docking resulted in most of TDF binding to a pocket region formed by its three stem loops (Figure 3B). While M2 docking showed a less uniform pattern of TDF binding sites with the density of sites spread in the central region of the structure (Figure 3C, Supplementary Data 4).

### 3.3 Interaction parameters between M1/M2 and TDF

Our M1 and M2 aptamers were chemically synthesized and the binding to the TDF was assayed using two different approaches: pulldown assays using unlabeled aptamers to test specificity and the Stopped flow (SF) technique to monitor the kinetics of binding. For the pulldown assays, we prepared seven reactions containing 100 nM of the TDF protein or a control protein immobilized to magnetic beads, and 300 nM of M1, M2 or a control oligonucleotide. After the incubation and magnetic separation, the protein-oligonucleotide complexes were resuspended in 20 μL of SELEX buffer 1X and incubated at 95°C for 3 min to allow dissociation of the aptamers. Then, the supernatants were collected and measured at 260 nm using a Nanodrop One spectrophotometer (Thermo Fisher Scientific, USA) in order to determine the concentration of bound oligonucleotides. M1 and M2 aptamers showed being more specific for TDF than for PKnG kinase domain. Consistently, binding of TDF to an unrelated oligonucleotide was lower than M1 or M2, yet we could not find a significant difference, likely due to the large standard deviation of the mean (Figure 4).

**Figure 4.**
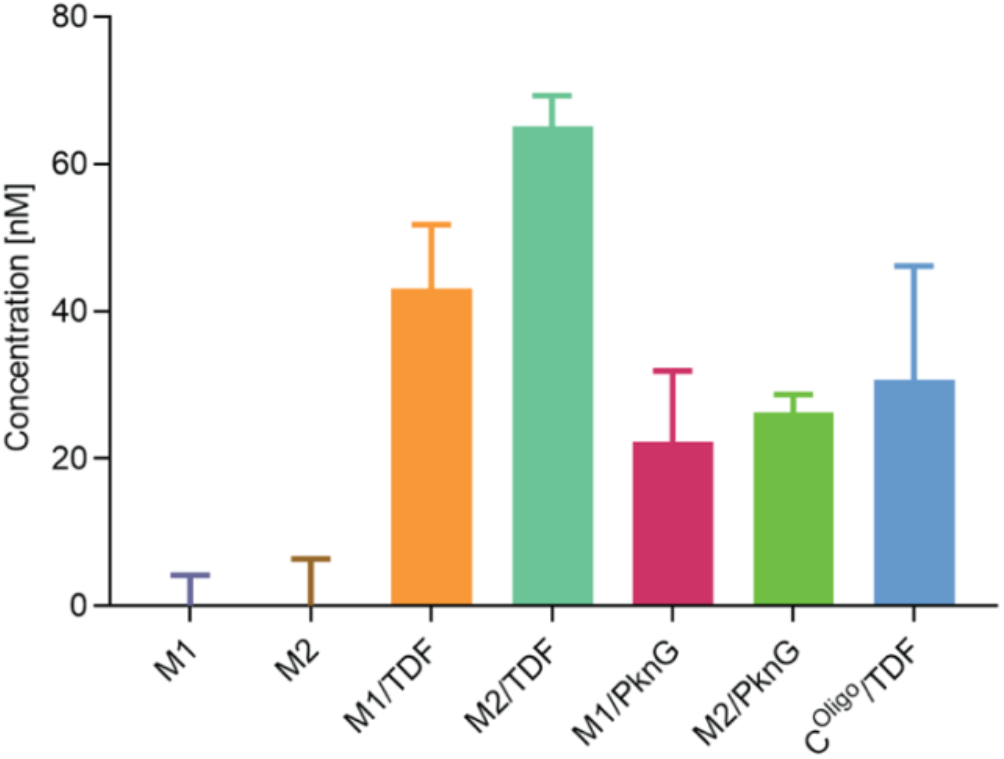
M1 and M2 aptamers bind preferentially to TDF. TDF and the non-related PKnG were subjected to interact with the randomized RNA library or with our M1 or M2 aptamers, then we performed a pulldown. The chart shows the concentration of the oligos that were retrieved after a pulldown using magnetic beads that were prebound to TDF or PKnG (unrelated protein control).

In order to further explore the mechanism of interaction between the aptamers and TDF, we measured the binding kinetics using the Stopped-Flow (SF) technique coupled to Förster Resonance Energy Transfer (FRET) between complementary probes to the 5’ and 3’ ends of our aptamers. The probes were 13 nucleotides long and were unmodified or with modifications: 5’ Fluorescein amidite (FAM) or 3’ Black Hole Quencher^®^ 1 (BHQ1) (Supplementary Table 2). The probes were annealed to our aptamers (see 2.7 Oligonucleotides synthesis and annealing) to see if the FAM-aptamer-BHQ1 complex (aptamer complex hereinafter) could be used as a beacon in the presence of the TDF (Figure 5A). Two complexes could be tested: FAM and BHQ1 modifications aiming inside and aiming outside, in and out configuration, respectively (Figure 5A). We mixed the pre-annealed aptamers against a molar excess of TDF in the SF to determine the effectiveness of the two probes configurations (In and Out). The In configuration resulted in a greater change in fluorescence for both M1 and M2 (Supplementary Figure 4).

**Figure 5.**
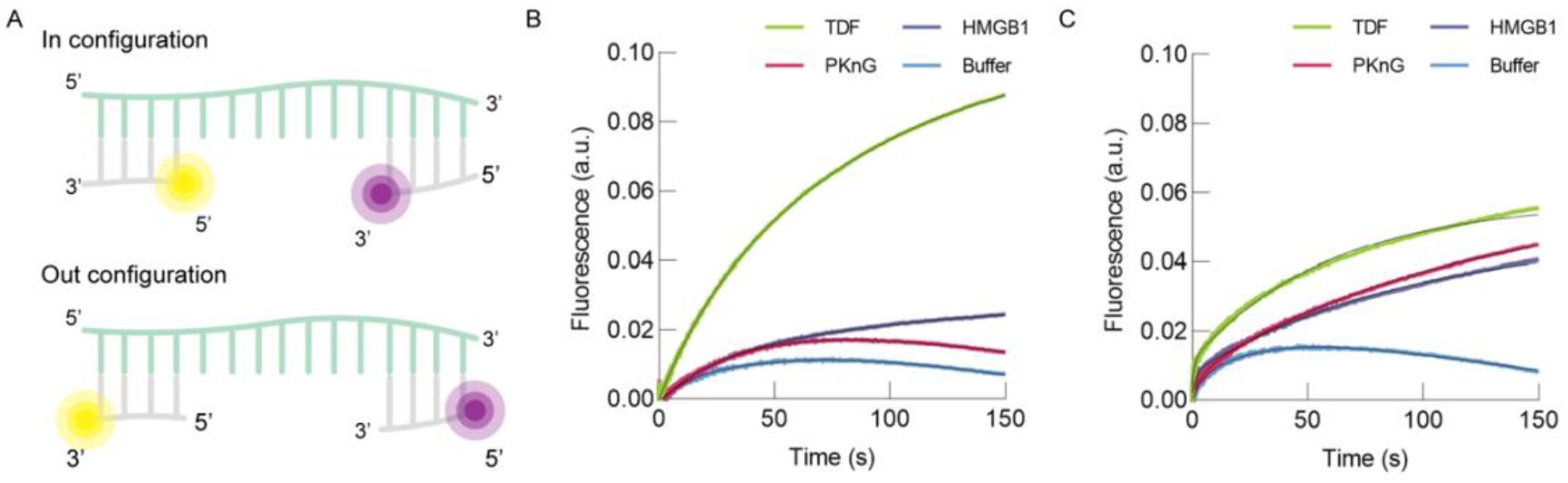
M1 rapidly interacts with TDF. (A) Scheme of the aptamer complex formed by the interaction of M1 or M2 aptamer (green) with FAM-modified probes (gray and yellow) and BHQ1-modified probes (gray and purple) aiming inside (top) and outside (bottom) the aptamer. The best signal was obtained when both probes (FAM and BHQ1) were aiming inside (Supplementary Figure 4). In order to determine the specificity and kinetics of our aptamers, the interaction of 100 nM of M1 (B) and M2 (C) were measured upon mixing with 250 nM TDF (green), HMGB1 (grey), PKnG (purple), or against buffer (light blue).

To verify that the change of fluorescence was due to an alteration in the distance between the FAM and BHQ1 modifications, we monitored fluorescence changes in rapid kinetic reactions between the probes. Mixing a preincubated FAM-aptamer complex against the BHQ1 probe resulted in a decrease in fluorescence, that results from diminishing the distance between fluorophores. While the interaction of FAM probe against BHQ1 probe or BHQ1 and FAM probes vs. TDF, or FAM-aptamer complex vs. non-modified probe resulted in no change in fluorescence (Supplementary Figure 5).

Regarding the specificity of our M1 and M2 aptamers to the TDF in the SF apparatus, aptamer complexes were rapidly mixed against different proteins: PKnG from *M. tuberculosis* and the HMG-box domain of the human HMGB1 (hereinafter called HMGB1). Mixing the FAM-M1-BHQ1 complex (M1 complex hereinafter) against the TDF resulted in an increase in fluorescence and there were no fluorescence changes upon mixing the complex against HMGB1 or PKnG (Figure 5B). The FAM-M2-BHQ1 complex (M2 complex hereinafter) against the TDF had a similar pattern as the M1 complex, but with a short amplitude. The M2 complex against HMGB1 or PKnG resulted in an increase in fluorescence of slightly less intensity than TDF (Figure 5C).

## 4. Discussion

In this study, we show that the aptamer beacon M1 binds specifically to TDF and provides a fluorescence signal that could be readily measured in fluorescence plate readers, unlocking a number of potential uses to localize TDF. To our knowledge, M1 is the first aptamer reported for TDF. Aptamers has been mainly used as biomarker detectors (Hamaguchi, Ellington, and Stanton 2001; Liang et al. 2011). Their principal strength comes from its great versatility of being modified with different chemical groups (Gawande et al. 2017; Hamaguchi, Ellington, and Stanton 2001). In this work, we show that our M1 and M2 aptamers show a degree of specificity for its target TDF, and when modified allowed to operate as beacons to monitor the binding to TDF. This approach has been previously used for the visualization endogenous proteins in living cells (Liang et al. 2011; Wang, Ding, and Zhou 2019). Theoretically, only small molecules with certain characteristics are able to penetrate tissues. However, it has been reported that even aptamers (~23.5 KDa) are able to penetrate tissues or even being translocated to the cells (Lenn et al. 2018; Wang, Ding, and Zhou 2019). Our M1 aptamer could be further tested for cell permeability in appropriate laboratory facilities that could unlock novel approaches for *in vivo* visualization of TDF in the cell.

The *SRY* gene and its protein product TDF has been of research interest since the discovery of its function in sexual differentiation (Wilhelm, Palmer, and Koopman 2007). The principal role of TDF during the embryo development is to work as a transcription factor for the expression of the Sox9 gene and the initiation of male differentiation (Sekido and Lovell-Badge 2009). However, its expression seems to go beyond the embryo development as it has been reported to be localized in midbrain dopamine neurons and regulates components of the synthesis of catecholamines (Czech et al. 2012). Also, its out-of-time activation could lead to a male-specific hepatocarcinogenesis due to its downstream targets (Liu et al. 2017). It is clear that its localization and function are relevant for the correct metabolism of male organisms, which puts TDF as an interesting target protein.

Regarding the area of *in vitro* fertilization, sperm sex sorting has been used to help prevent Duchene Muscular Dystrophy or hemophilia, which are sex-associated heritable diseases (Johnson et al. 1993). In this case, the difference between the size of the X and Y chromosomes in spermatozoa is detected by the Hoechst 33342 stain using a flow cytometer (Johnson et al. 1993; Sumner and Robinson 1976). However, the principal issue with this technique is the possibility of decreasing the sperm DNA integrity caused by the dye (Katigbak et al. 2019). Under this scenario, TDF and TDF-binding aptamer beacons could be used for sex selection. Although the localization of TDF in human ejaculated spermatozoa is yet unclear, round spermatids are already haploid cells and TDF has been reported to be present in them (Salas-Cortés et al. 1999, 2001).

Aptamers seem a valid alternative for the detection of different molecules of interest. The present study proposes potential new aptamer beacons as biosensors for the identification of the human TDF. These aptamer beacons can be further modified in order to develop alternative methods such as drug delivery or sex sorting.

## Author contributions

DFJ conceived the project. DJF, JANK, and PM obtained funding. DFJ and JANK performed experiments. DFJ, JANK, and PM analyzed the data. DFJ, JANK, and PM wrote the manuscript.

## Acknowledgments

We are very thankful to Catherine Paola Salvatierra Castro for her expert support on the artwork and graphics.

## Funding

This work was funded by the VI Annual Research Incentive Competition (EXP-19) from Universidad Peruana de Ciencias Aplicadas to DFJ, and grants from the Fondecyt [136-2016-Fondecyt (http://www.fondecyt.gob.pe/)] and InnovatePeru [297-INNOVATEPERU-EC-2016 (http://www.innovateperu.gob.pe/)] to PM.

## Competing Interests

The authors declare no competing interests.

## List of Supporting Information

**Supplementary Figure 1.**
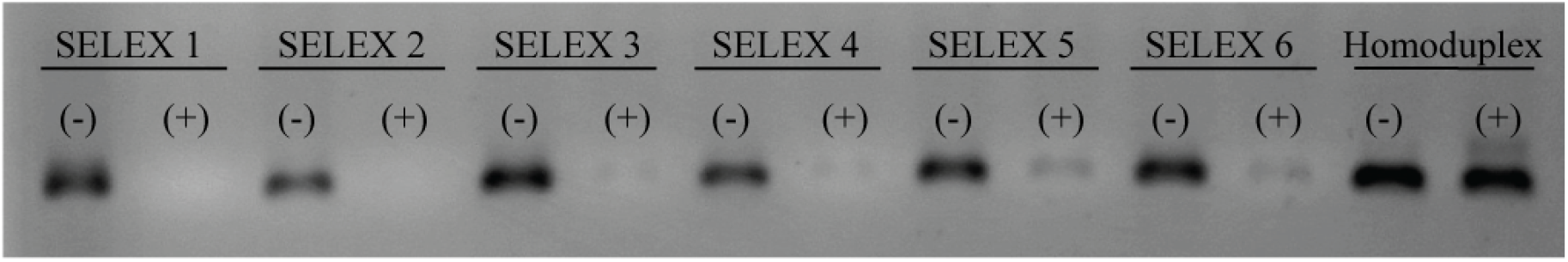
DiVE assay of the six SELEX cycles and a positive control (Homoduplex) with (+) or without (-) S1 endonuclease.

**Supplementary Figure 2.**
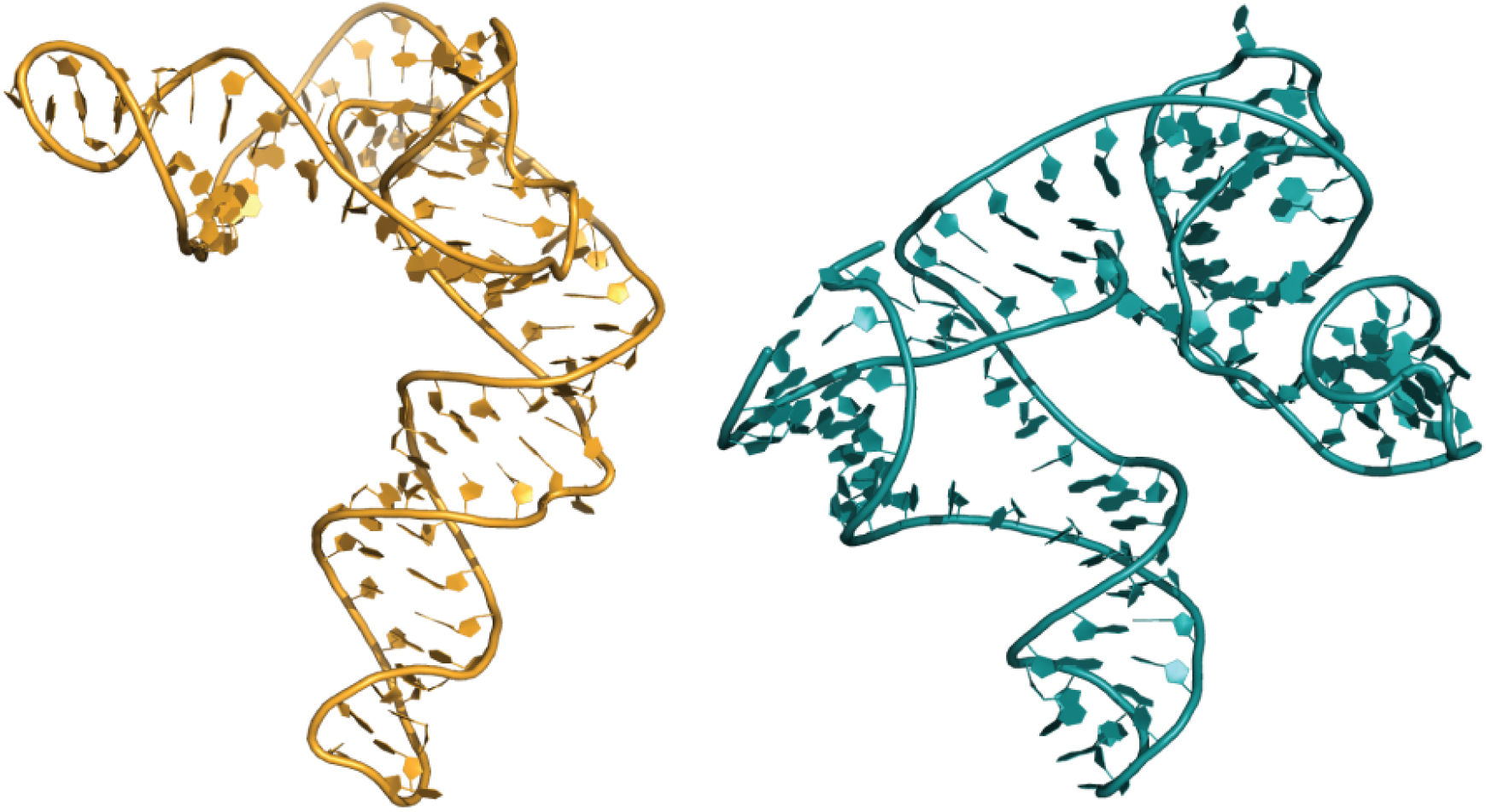
3D structures of M1 (left) and M2 (right) aptamers.

**Supplementary Figure 3.**
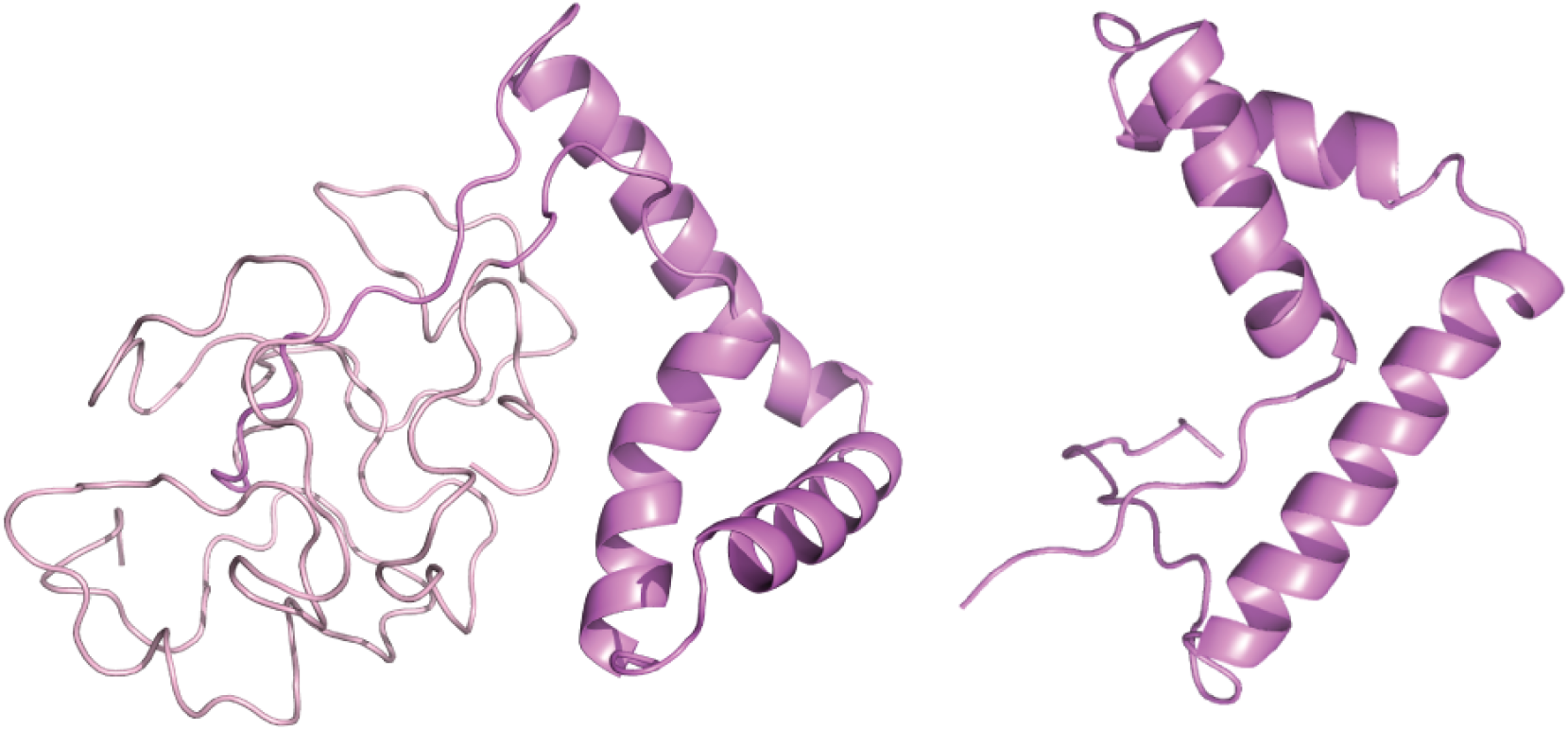
3D structures of the complete human TDF (left) and its HMG-box domain (right).

**Supplementary Figure 4.**
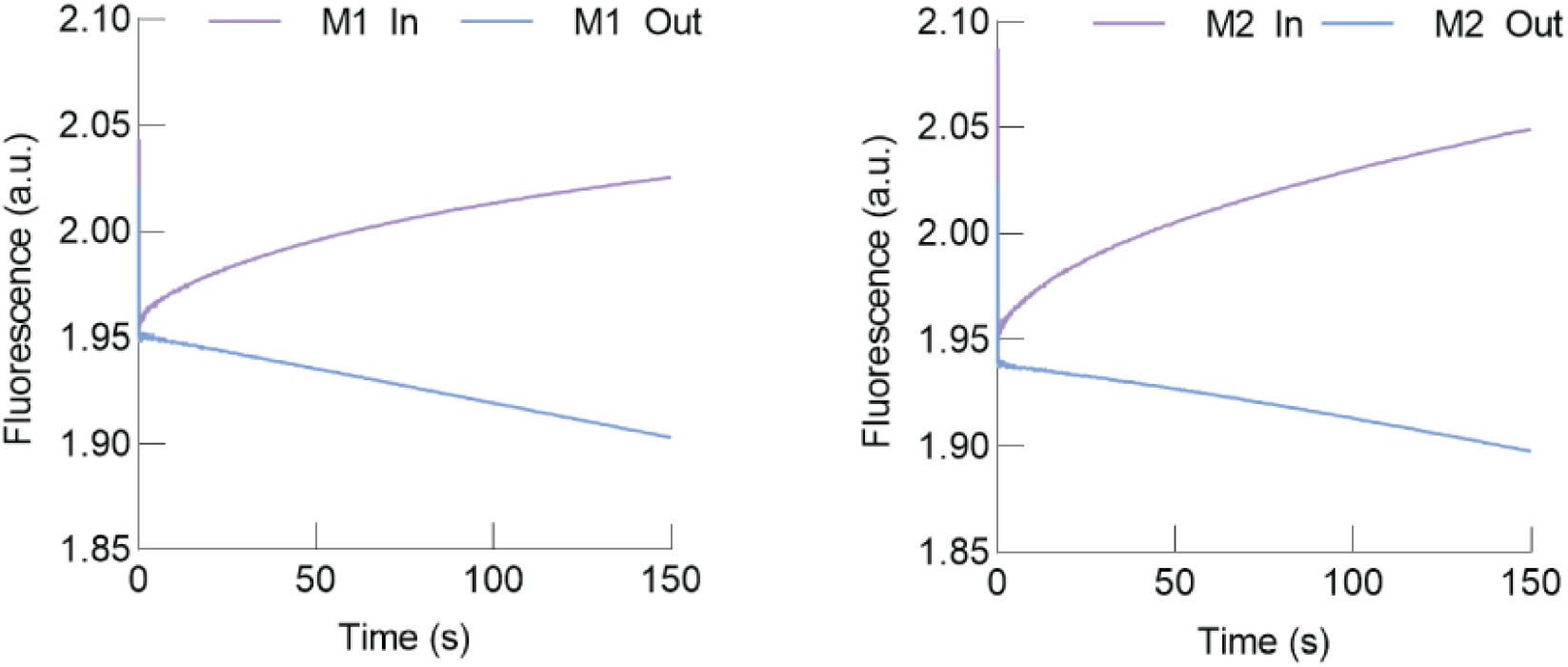
M1 (left) and M2 (right) In and Out complexes against TDF.

**Supplementary Figure 5.**
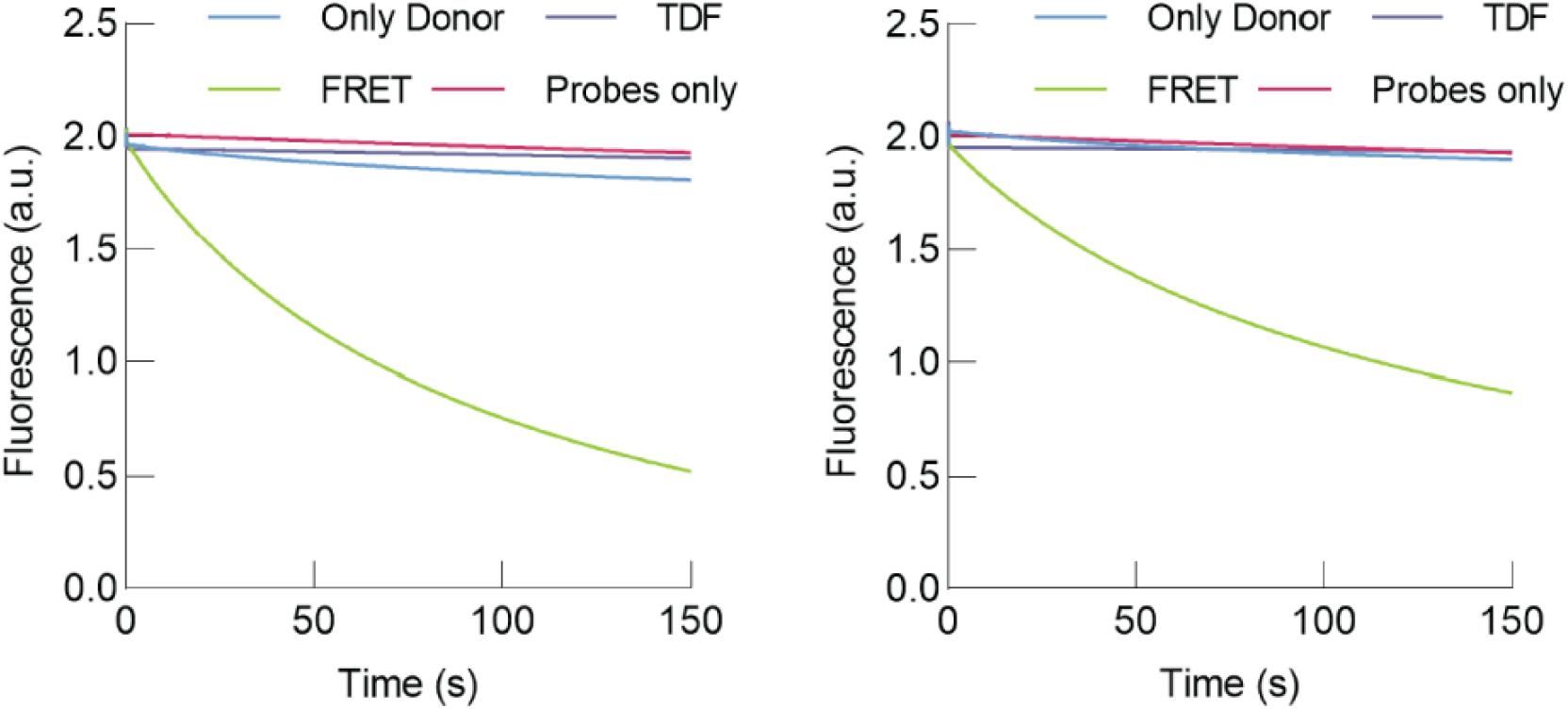
Fluorescence changes in rapid kinetic reactions for M1 (left) and M2 (right). *Only donor*: FAM-aptamer vs. non modified probe. *TDF*: FAM and BHQ1 probes vs. TDF. *FRET*: FAM-aptamer complex vs. BHQ1 probe. *Probes only*: FAM probe vs. BHQ1 probe.

**Supplementary Table 1.** Reactants and products quantities and concentrations, as well as conditions in each SELEX cycle.

| *SELEX cycle* | *1* | *2* | *3* | *4* | *5* | *6* |
| --- | --- | --- | --- | --- | --- | --- |
| RNA (pmol) | 1000 | 100 | 100 | 75 | 75 | 75 |
| TDF (pmol) | 100 | 50 | 25 | 15 | 15 | 15 |
| Final volume (μL) | 200 | 200 | 300 | 500 | 1500 | 1500 |
| RNA concentration (μM) | 5 | 0.5 | 0.3 | 0.15 | 0.05 | 0.05 |
| TDF concentration (μM) | 0.5 | 0.25 | 0.08 | 0.03 | 0.01 | 0.01 |
| Incubation (min) | 60 | 20 | 20 | 15 | 5 | 5 |
| Washes (#) | 0 | 3 | 3 | 5 | 5 | 5 |
| RT-PCR (PCR cycles) | 6 | 5 | 5 | 7 | 6 | 6 |
| RT-PCR product: dsDNA (pmol) | 162.45 | 284.21 | 179.45 | 145.14 | 143.02 | 110.83 |
| Transcription (min) | 120 min |  |  |  |  | -- |
| Transcription product: RNA (pmol) | 113.19 | 847.77 | 450.36 | 726.48 | 454.57 | -- |

**Supplementary Table 2.**
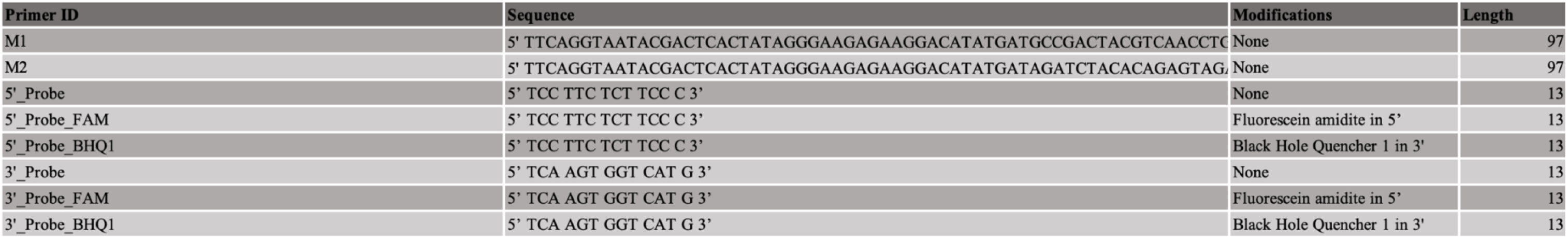
List of oligonucleotides (DNA) used for the pull down and stopped flow analysis. M1 and M2 oligonucleotides were synthesized as duplexes.

**Supplementary Data 1.** Aptamer frequency analysis by the FASTAptamer-Count algorithm.

**Supplementary Data 2.** Aptamer ranking of all SELEX cycles.

**Supplementary Data 3.** Aptamer enrichment analysis by the FASTAptamer-Enrich algorithm.

**Supplementary Data 4.** Top 100 MOE S-scores for M1 and M2 aptamers.

## Notes

### Competing Interest Statement

The authors have declared no competing interest.

## References

Afgan, Enis et al. 2018. “The Galaxy Platform for Accessible, Reproducible and Collaborative Biomedical Analyses: 2018 Update.”Nucleic Acids Research 46(W1): W537–44.

Alam, Khalid K., Jonathan L. Chang, and Donald H. Burke. 2015. “FASTAptamer: A Bioinformatic Toolkit for High-Throughput Sequence Analysis of Combinatorial Selections.”Molecular Therapy - Nucleic Acids 4(3): 1–10.

Antczak, Maciej et al. 2016. “New Functionality of RNAComposer: An Application to Shape the Axis of MiR160 Precursor Structure.”Acta Biochimica Polonica 63(4): 737–44.

Bailey, Timothy L. et al. 2009. “MEME Suite: Tools for Motif Discovery and Searching.”Nucleic Acids Research 37(SUPPL. 2): 202–8.

Chulluncuy, Roberto et al. 2016. “Conformational Response of 30S-Bound IF3 to A-Site Binders Streptomycin and Kanamycin.”Antibiotics 5(4): 1–14.

Clepet, Christian et al. 1993. “The Human SRY Transcript.”Human Molecular Genetics 2(12): 2007–12.

Czech, Daniel P. et al. 2012. “The Human Testis-Determining Factor SRY Localizes in Midbrain Dopamine Neurons and Regulates Multiple Components of Catecholamine Synthesis and Metabolism.”Journal of Neurochemistry 122(2): 260–71.

Denny, P., S. Swift, F. Connor, and A. Ashworth. 1992. “An SRY-Related Gene Expressed during Spermatogenesis in the Mouse Encodes a Sequence-Specific DNA-Binding Protein.”EMBO Journal 11(10): 3705–12.

Ellis, Peter James, and Robert P Erickson. 2017. “149 - Genetics of Sex Determination and Differentiation.” In eds. Richard A Polin et al. Elsevier, 1510–1519.e4.

Gawande, Bharat N. et al. 2017. “Selection of DNA Aptamers with Two Modified Bases.”Proceedings of the National Academy of Sciences 114(11): 2898–2903.

George, Fredrick W, and Jean D Wilson. 1984. “2 - Sexual Differentiation.” In eds. RICHARD W BEARD and PETER W B T - Fetal Physiology and Medicine (Second Edition) NATHANIELSZ. Butterworth-Heinemann, 57–79.

Goodfellow, P. N., and S. M. Darling. 1988. “Genetics of Sex Determination in Man and Mouse.”Development 102(2): 251–58.

Gruber, Andreas R. et al. 2008. “The Vienna RNA Websuite.”Nucleic acids research 36(Web Server issue): 70–74.

Hamaguchi, Nobuko, Andrew Ellington, and Martin Stanton. 2001. “Aptamer Beacons for the Direct Detection of Proteins.”Analytical Biochemistry 294(2): 126–31.

Harley, Vincent R., Michael J. Clarkson, and Anthony Argentaro. 2003. “The Molecular Action and Regulation of the Testis-Determining Factors, SRY (Sex-Determining Region on the Y Chromosome) and Sox9 [Sry-Related High-Mobility Group (HMG) Box 9].”Endocrine Reviews 24(4): 466–87.

Illumina. 2013. “16S Metagenomic Sequencing Library.”Illumina.com (B): 1–28.

Johnson, Lawrence A. et al. 1993. “Gender Preselection in Humans? Flow Cytometric Separation of x and y Spermatozoa for the Prevention of x-Linked Diseases.”Obstetrical and Gynecological Survey 49(5): 342–44.

Joseph, Diego F. et al. 2019. “DNA Aptamers for the Recognition of HMGB1 from Plasmodium Falciparum.”PLoS ONE 14(4): 1–20.

Kanai, Yoshiakira, Ryuji Hiramatsu, Shogo Matoba, and Tomohide Kidokoro. 2005. “From SRY to SOX9: Mammalian Testis Differentiation.”Journal of Biochemistry 138(1): 13–19.

Katigbak, Roberto D. et al. 2019. “Review on Sperm Sorting Technologies and Sperm Properties toward New Separation Methods via the Interface of Biochemistry and Material Science.”Advanced Biosystems 3(9): 1–16.

Kelley, Lawrence A et al. 2016. “The Phyre2 Web Portal for Protein Modeling, Prediction and Analysis.”Nature Protocols 10(6): 845–58.

Kiefer, Julie C. 2007. “Back to Basics: Sox Genes.”Developmental Dynamics 236(8): 2356–66.

Kodrič, Klemen et al. 2019. “Sex-Determining Region Y (SRY) Attributes to Gender Differences in RANKL Expression and Incidence of Osteoporosis.”Experimental and Molecular Medicine 51(8).

Lenn, Jon D. et al. 2018. “RNA Aptamer Delivery through Intact Human Skin.”Journal of Investigative Dermatology 138(2): 282–90.

Li, Chunjin et al. 2011. “Detection of the SRY Transcript and Protein in Bovine Ejaculated Spermatozoa.”Asian-Australasian Journal of Animal Sciences 24(10): 1358–64.

Liang, Yu et al. 2011. “Aptamer Beacons for Visualization of Endogenous Protein HIV-1 Reverse Transcriptase in Living Cells.”Biosensors and Bioelectronics 28(1): 270–76.

Lim, Theam Soon et al. 2011. “Diversity Visualization by Endonuclease: A Rapid Assay to Monitor Diverse Nucleotide Libraries.”Analytical Biochemistry 411(1): 16–21.

Liu, Chang et al. 2017. “Activation of SRY Accounts for Male-Specific Hepatocarcinogenesis: 18 Implication in Gender Disparity of Hepatocellular Carcinoma.”Cancer Letters 410(2017): 20–31.

Milon, Pohl, Andrey L. Konevega, Claudio O. Gualerzi, and Marina V. Rodnina. 2008. “Kinetic Checkpoint at a Late Step in Translation Initiation.”Molecular Cell 30(6): 712–20.

Modi, D. et al. 2005. “Ontogeny and Cellular Localization of SRY Transcripts in the Human Testes and Its Detection in Spermatozoa.”Reproduction 130(5): 603–13.

Molecular Operating Environment (MOE), 2019.01; Chemical Computing Group ULC, 1010 Sherbooke St. West, Suite #910, Montreal, QC, Canada, H3A 2R7, 2020.

Olivares, Aleida et al. 2018. “Regulation of CATSPER1 Expression by the Testis-Determining Gene SRY.”PLoS ONE 13(10): 1–16.

Popenda, Mariusz et al. 2012. “Automated 3D Structure Composition for Large RNAs.”Nucleic Acids Research 40(14): 1–12.

Poulat, Francis et al. 1995. “Nuclear Localization of the Testis Determining Gene Product SRY.”Journal of Cell Biology 128(5): 737–48.

R. Oliphant, Arnold, Christopher J. Brandl, and Kevin Struhl. 1989. “Defining the Sequence Specificity of DNA-Binding Proteins by Selecting Binding Sites from Random-Sequence Oligonucleotides: Analysis of Yeast GCN4 Protein.”Molecular and cellular biology 9(7): 2944–49.

Salas-Cortés, Laura et al. 1999. “The Human SRY Protein Is Present in Fetal and Adult Sertoli Cells and Germ Cells.”International Journal of Developmental Biology 43(2): 135–40.

Salas-Cortés, Laura et al. 2001. “Expression of the Human SRY Protein during Development in Normal Male Gonadal and Sex-Reversed Tissues.”Journal of Experimental Zoology 290(6): 607–15.

Schneider, Caroline A, Wayne S Rasband, and Kevin W Eliceiri. 2012. “NIH Image to ImageJ: 25 Years of Image Analysis.”Nature Methods 9: 671. https://doi.org/10.1038/nmeth.2089.

Sekido, Ryohei, and Robin Lovell-Badge. 2009. “Sex Determination and SRY: Down to a Wink and a Nudge?”Trends in Genetics 25(1): 19–29.

Sinclair, Andrew H. et al. 1990. “A Gene from the Human Sex-Determining Region Encodes a Protein with Homology to a Conserved DNA-Binding Motif.”Nature 346(6281): 240–44.

Stoltenburg, R., C. Reinemann, and B. Strehlitz. 2005. “FluMag-SELEX as an Advantageous Method for DNA Aptamer Selection.”Analytical and Bioanalytical Chemistry 383(1): 83–91.

Su, H., and Y. F.C. Lau. 1993. “Identification of the Transcriptional Unit, Structural Organization, and Promoter Sequence of the Human Sex-Determining Region Y (SRY) Gene, Using a Reverse Genetic Approach.”American Journal of Human Genetics 52(1): 24–38.

Sumner, A. T., and J. A. Robinson. 1976. “A Difference in Dry Mass between the Heads of X and Y Bearing Human Spermatozoa.”Journal of Reproduction and Fertility 48(1): 9–15.

Thiviyanathan, Varatharasa, and David G. Gorenstein. 2012. “Aptamers and the next Generation of Diagnostic Reagents.”Proteomics - Clinical Applications 6(11–12): 563–73.

Tricoli, James V., Joyce L. Yao, Sharon A. D’Souza, and R. Bruce Bracken. 1993. “Detection of Sex-region Y (SRY) Transcripts in Human Prostate Adenocarcinoma and Benign Prostatic Hypertrophy.”Genes, Chromosomes and Cancer 8(1): 28–33.

Tuerk, C, and L Gold. 1990. “Systematic Evolution of Ligands by Exponential Enrichment: RNA Ligands to Bacteriophage T4 DNA Polymerase.”Science (New York, N.Y.) 249(4968): 505–10.

Wang, Shengfeng, Jinsong Ding, and Wenhu Zhou. 2019. “An Aptamer-Tethered, DNAzyme-Embedded Molecular Beacon for Simultaneous Detection and Regulation of Tumor-Related Genes in Living Cells.”Analyst 144(17): 5098–5107.

Waterhouse, Andrew et al. 2018. “SWISS-MODEL: Homology Modelling of Protein Structures and Complexes.”Nucleic Acids Research 46(W1): W296–303.

Wilhelm, Dagmar, Stephen Palmer, and Peter Koopman. 2007. “Sex Determination and Gonadal Development in Mammals.”Physiological Reviews 87(1): 1–28.

Wu, Yi Xi, and Young Jik Kwon. 2016. “Aptamers: The ‘Evolution’ of SELEX.”Methods 106: 21–28.

Zuker, Michael. 2003. “Mfold Web Server for Nucleic Acid Folding and Hybridization Prediction.”Nucleic Acids Research 31(13): 3406–15.

